# Adipocyte-specific modulation of KLF14 expression in mice leads to sex-dependent impacts in adiposity and lipid metabolism

**DOI:** 10.1101/2021.09.15.460489

**Authors:** Qianyi Yang, Jameson Hinkle, Jordan N. Reed, Redouane Aherrahrou, Zhiwen Xu, Thurl E. Harris, Erin J. Stephenson, Kiran Musunuru, Susanna R. Keller, Mete Civelek

## Abstract

Genome-wide association studies identified single nucleotide polymorphisms on chromosome 7 upstream of *KLF14* to be associated with metabolic syndrome traits and increased risk for Type 2 Diabetes (T2D). The associations were more significant in women than in men. The risk allele carriers expressed lower levels of the transcription factor KLF14 in adipose tissues than non-risk allele carriers. To investigate how adipocyte KLF14 regulates metabolic traits in a sex-dependent manner, we characterized high-fat diet fed male and female mice with adipocyte-specific *Klf14* deletion or overexpression. *Klf14* deletion resulted in increased fat mass in female mice and decreased fat mass in male mice. Female *Klf14-*deficient mice had overall smaller adipocytes in subcutaneous fat depots but larger adipocytes in parametrial depots, indicating a shift in lipid storage from subcutaneous to visceral fat depots. They had reduced metabolic rates and increased respiratory exchange ratios consistent with increased utilization of carbohydrates as an energy source. Fasting and isoproterenol-induced adipocyte lipolysis was defective in female *Klf14-*deficient mice and concomitantly adipocyte triglycerides lipase mRNA levels were downregulated. Female *Klf14-*deficient mice cleared blood triglyceride and NEFA less efficiently than wild type. Finally, adipocyte-specific overexpression of *Klf14* resulted in lower total body fat in female but not male mice. Taken together, consistent with human studies, adipocyte KLF14 deficiency in female but not in male mice causes increased adiposity and redistribution of lipid storage from subcutaneous to visceral adipose tissues. Increasing KLF14 abundance in adipocytes of females with obesity and T2D may provide a novel treatment option to alleviate metabolic abnormalities.

## INTRODUCTION

Genome-Wide Association Studies (GWAS) identified several genetic variants on chromosome 7 upstream of the *KLF14* gene to be associated with a multitude of metabolic abnormalities, including insulin resistance, type 2 diabetes, and coronary artery disease (1–4). The associations were more pronounced in women than in men (5). GWAS of gene expression performed in the Multiple Tissue Human Expression Resource (MuTHER) and the Metabolic Syndrome in Men (METSIM) cohorts identified KLF14 as a master regulator of gene expression in subcutaneous adipose tissue, despite its nearly ubiquitously expression across all human tissues (6–9). GWAS-associated single nucleotide polymorphisms (SNPs) were also associated with KLF14 expression levels in *cis* and nearly 400 genes in *trans* in adipose tissue (1,6,10).The locus is one of the largest *trans*-eQTL hotspots known in the human genome (7). The results also suggested that disease-associated variants act in adipose tissue to increase disease risk. However, given that adipose tissue contains a multitude of cell types, it was not clear which cells were ultimately responsible for KLF14’s adverse effects.

Several studies have implicated KLF14, a single exon gene in the Krüpple-like factor family of transcription factors, in the development of metabolic diseases, including obesity, insulin resistance, and T2D. Small *et al*. demonstrated that lowering KLF14 expression in preadipocytes from females prevented maturation to adipocytes as measured by the expression of adipogenesis markers and lipid accumulation (1). Histological analysis of subcutaneous adipose tissue biopsies showed that females who were homozygous for the T2D risk allele had larger adipocytes, suggesting that lower KLF14 expression is associated with adipocyte dysfunction (1). Collectively, these studies implied that KLF14 plays roles in adipogenesis and mature adipocyte size and function. T2D-associated variants also have strong associations with high density lipoprotein and triglyceride levels in the serum (11,12). However, the mechanisms by which KLF14 regulates adipocyte differentiation and function are not known.

To further investigate the role of KLF14 in adipocytes and to establish a mouse model that is relevant to humans, we characterized mice with adipocyte-specific *Klf14* deletion or overexpression. Our findings demonstrate that adipocyte KLF14 is a key regulator of lipid metabolism and, consistent with findings in human, this regulation is female-specific. Our findings may facilitate the development of novel therapeutics to treat obesity and related metabolic diseases specifically in females.

## MATERIALS and METHODS

### Generation of mice with adipocyte-specific deletion and overexpression of *Klf14*

The generation of adipocyte-specific *Klf14* knockout mice has been described in detail elsewhere (1). Briefly, *in vitro* transcribed Cas9 mRNA, two gRNAs designed to target sites upstream and downstream of the *Klf14* gene locus, two single-strand DNA oligonucleotides bearing loxP sequences along with 80-nt homology arms matching the target sites were injected into the cytoplasm of fertilized oocytes from C57BL/6J mice (Supplementary Figure. 1A). Genomic DNA samples from founders were screened for correct loxP sequences flanking the *Klf14* gene by PCR and confirmed by Sanger sequencing. Mice with two loxP sites that segregated on the same chromosome were bred with *Adipoq*-Cre mice on the C57BL/6J background [B6; FVB-Tg (*Adipoq*-Cre)1Evdr/J, The Jackson Laboratory] to obtain litters with homozygous *Klf14* loxP knock-in mice that were Cre+ (*Klf14*^fl/fl^*Adipoq*-Cre+ or KO) and wild type mice that were positive for *Klf14* loxP but negative for *Adipoq*-Cre (*Klf14*^fl/fl^*Adipoq*-Cre- or WT) for experiments. The presence of the *Adipoq*-Cre allele was confirmed by PCR (Supplementary Table 1). Absence of KLF14 protein expression in adipocytes isolated from knockout animals was confirmed by Western Blotting (Supplementary Figure 1B). To generate mice with adipocyte-specific overexpression (OE) of *Klf14, Klf14* was PCR amplified from C57BL/6J cDNA and cloned downstream of the 5.43 Kb Adiponectin (*Adipoq*) promoter at an EcoRV site using Gibson Assembly into the pADNpcDNA3.1 KanR vector. The 8.2 Kb expression cassette containing the *Adipoq* promoter, *Klf14* cDNA, and 3’ UTR was gel purified after restriction digestion with NotI and NgoMIV. The transgene was injected into pronuclei of fertilized C57BL/6N eggs to produce random integrants at the University of California, Irvine Transgenic Mouse Facility. The presence of the transgene was confirmed by PCR (Primers are listed in Supplementary Table 1). We labeled these mice *Adipoq-Klf14-*OE. Increased KLF14 protein expression in adipocytes isolated from transgenic animals was confirmed by Western Blotting (Supplementary Figure 1C).

### Animal Husbandry

All animal protocols were approved by the University of Virginia Animal Care and Use Committee. Mice were maintained in a 12-hour light/12-hour dark cycle with free access to water and standard chow (Envigo Teklad LM-485 irradiated mouse/rat sterilizable diet. Cat No. 7912). For high fat diet (HFD) studies, the mice were fed a rodent diet with 40% Kcal fat, 43% Kcal carbohydrate and 17% Kcal protein (Research Diets, Inc., Cat No. D12079B). The mice were euthanized by cervical dislocation after induction of deep anesthesia using isoflurane, consistent with the recommendations of the Panel of the American Veterinary Medical Association.

### Body composition analysis

Total body fat and lean mass were assessed in conscious mice using a noninvasive quantitative magnetic resonance imaging system (Echo Medical Systems, Cat No. EchoMRI™ 3-in-1 v2.1;), as previously reported (13).

### Glucose and insulin tolerance tests

Glucose and insulin tolerance tests were performed as previously described (14). For the glucose tolerance test, mice were fasted from 7 AM to 1 PM and were administered glucose at 1 mg/g body weight intraperitoneally. Blood glucose levels were measured with a OneTouch Ultra Glucometer (Lifescan) in blood drops obtained from nicked tail veins before and at 10, 20, 30, 60, 90 and 120 min after the injection. The insulin tolerance test was performed at 1–2 PM on mice fed ad libitum. Insulin (Humulin, 100 U/ml, from Eli Lilly) at 0.75 U/kg body weight was administered by intraperitoneal injection of a 0.25 U/ml solution in 0.9% NaCl. Blood glucose levels were measured immediately before and at 15, 30, 45, 60, and 90 min after the injection.

### Plasma insulin and lipid analysis

Mice were either fed ad libitum, and samples taken between 7 AM and 8 AM, or they were fasted for 6 hours from 7 AM to 1 PM before samples were taken. Whether measurements were done in the fed or fasted state is indicated in the Results section. For insulin, fatty acids, and glycerol measurements, blood samples were collected either through retro-orbital bleeding or cardiac puncture. For insulin, high density lipoprotein (HDL), total cholesterol, and triglyceride (TG) measurements, plasma was obtained from the blood by the addition of heparin at 10 U/ml, followed by centrifugation at 7800 rcf in a microfuge for 10 min at 4°C. Insulin levels were measured with an ELISA (Alpco, Cat No. 80-INSMSU-E01). Total cholesterol, HDL, and TG levels were measured using colorimetric assays (FUJIFILM Wako Diagnostics, Cat Nos. 999-02601, 997-72591, 464-01601).

### Indirect calorimetry analysis

For determination of energy expenditure and respiratory exchange ratios, mice were placed in Oxymax metabolic chambers (Columbus Instruments), under a constant environmental temperature (22 °C) and a 12-hour light, 12-hour dark cycle as previously described (15). Oxygen consumption (V_O2_), carbon dioxide production (V_CO2_), and ambulatory activity were determined for each mouse every 15 min over a 72-h period. Experimental mice were acclimated to the cages during the first 24 hours. Measurements taken during the following 48 hours were used for data analysis. Respiratory exchange ratios (RER) were calculated as carbon dioxide production/oxygen consumption (V_CO2_/V_O2_). Carbohydrate utilization was calculated as (20 kJ/L × V_O2_ uptake) × [(RER - 0.7)/0.3] and fat utilization was calculated as (20kJ/ × V_O2_uptake) × (1 - [(RER × 0.7)/0.3)] (16). Both body fat and lean mass were included as covariates in the models used to analyze data obtained from the CLAMS experiments. Mice in each chamber had free access to water and food. Locomotor activity was monitored by a multidimensional infrared light beam system surrounding each cage.

### *In vivo* lipolysis

To determine lipolysis *in vivo*, two different experiments were performed. For lipolysis under fasting condition, blood samples were drawn from tail veins of 24-week-old mice using non-heparinized glass capillary tubes at 5 PM before food was withdrawn and 9 AM after 16-hour fasting to measure non-esterified fatty acids (NEFA) and glycerol. For β-adrenergic receptor stimulation experiments, blood samples were drawn from tail veins of random-fed 25-week-old mice via non-heparinized glass capillary tubes before stimulation. After a short recovery period (30 minutes), mice were injected intraperitoneally with 10 mg/kg isoproterenol (prepared in saline; Sigma Aldrich Cat No I6504). A second blood sample was collected 15 min post-injection. Blood from non-heparinized capillary tubes was allowed to clot on ice before centrifugation at 500 g for 20 min at 4 °C. Glycerol and NEFA concentrations in serum were determined colorimetrically using commercially available reagents (Sigma Aldrich, Cat No. F6428, Wako Life Sciences, Inc., Cat No. NEFA-HR (2)).

### *In vivo* lipid challenge

Experimental mice were fasted for 4 hours before the test (7 –11 AM). Olive oil (6 µl/g body weight) was given by oral gavage as described previously (17). Blood samples were drawn via heparinized (for TG)/non-heparinized (for NEFA) capillary tubes from tail veins before (time 0) and at 1, 2, 3, and 4 hours after gavage. TG and NEFA measurements were performed as described above.

### Determination of adipocyte sizes

Subcutaneous white adipose tissue (WAT) and parametrial-periovarian/epidydimal WAT from four mice of each sex and genotype were excised and fixed in 10% neutral buffered formalin (18). Tissues were embedded in paraffin, and 5 µm thick sections were prepared and stained with hematoxylin and eosin (H&E) at the University of Virginia Research Histology Core. Slides were scanned and images were acquired with an EVOS microscope camera. Quantification of adipocyte size was done using ImageJ (version 1.48) (19). Images were processed as described previously (20).

### Sex hormone measurements

Mouse sera collected at the time of euthanasia were stored at −80°C. After thawing, they were allowed to equilibrate at room temperature and spun to remove aggregates. 100 μL of 1:10 diluted sera per well were aliquoted for enzyme-linked immunosorbent assays (ELISA) in 96-well plates. Free testosterone, progesterone and estradiol levels were measured using Testosterone Parameter assay kit (R&D Systems, Cat No. KGE010), Progesterone ELISA Kit (Cayman Chemical, Cat No. 582601) and Estradiol ELISA Kit (Cayman Chemical, Cat No. 501890) following manufacturers’ instructions (21).

### Primary mouse adipocyte isolation

Adipocytes were isolated from subcutaneous and parametrial-periovarian/epididymal WAT as previously described (22). Immediately after euthanization fat pads were removed, placed in Krebs Ringer HEPES (KRH)– BSA buffer containing collagenase type I (Worthington Biochemical Corp., 1 mg/ml, 2 mg/g of tissue) and minced with scissors. Small tissue pieces were incubated in a 37°C shaking water bath (100 rpm) for 1 hour. Fat cells were separated from non-fat cells and undigested debris by filtration through a 0.4 mm Nitex nylon mesh (Tetko) and four washes by flotation with KRH buffer without BSA. After centrifugation at 1000 g for 15 minutes at 4°C the isolated fat cells floating on the surface were used for total RNA extraction and preparation of cell lysates for immunoblotting.

### Total RNA extraction, cDNA synthesis and real-time PCR

Total RNA was isolated from parametrial-periovarian/epididymal and subcutaneous adipocytes using a combination of Trizol Reagent® (Thermo fisher Scientific, USA) and miRNeasy kit (Qiagen, Germany) according to the manufacturers’ instructions (23). Adipocytes isolated from 200 mg of tissue in the previous step were homogenized in 2 ml of Trizol Reagent® using tissue-tearor homogenizer (Model 985370, BioSpec Products, Inc). After homogenization, samples were incubated at room temperature for 5 minutes and centrifuged at 12,000 g for 10 min at 4°C. The resulting fat monolayer on top was carefully avoided when pipetting the rest of the sample into a clean tube. 400 μl of chloroform was then added and samples mixed by vortexing. Samples were kept at room temperature for 3 minutes before centrifugation at 12,000 g for 30 min at 4°C. After centrifugation, samples separated into three phases with the RNA in the upper phase. The RNA phase was transferred to a new tube without disturbing the interphase. Sample volumes were measured and 1.5x sample volumes of 100% ethanol added. Samples were mixed thoroughly by inverting tubes several times. Samples were then loaded on miRNeasy spin columns (Qiagen) and the manufacturer’s protocol was followed for subsequent steps. Total RNA was eluted in 30 μl of RNAse-free water. Total RNA concentrations were quantified using a Qubit Fluorometer and a Qubit RNA BR assay kit (Thermo Fisher Scientific). 1 µg of total RNA was reverse transcribed using SuperScript IV reverse transcriptase (Invitrogen). Real-time PCR was performed in the QuantStudio™ 5 Real-Time PCR System (ThermoFisher Scientific) using SYBR Green Master Mix (Roche) and gene-specific primers (Supplementary Table 2).

### Protein extraction and Western blotting

Total protein was extracted from cells or tissues using cell lysis buffer (Cell Signaling, Cat No. 9803), containing 20 mM Tris-HCl (pH 7.5), 150 mM NaCl, 1 mM Na_2_EDTA, 1 mM EGTA, 1% Triton, 2.5 mM sodium pyrophosphate, 1 mM beta-glycerophosphate, 1 mM Na_3_VO_4_, and 1 µg/ml leupeptin. Protease inhibitor cocktail (ThermoFisher Scientific) and phosphatase inhibitor (ThermoFisher Scientific) were added to the lysis buffer before extraction. Protein concentrations were determined using a Pierce BCA Protein Assay Reagent (Pierce Biotechnology). Proteins (10 ug protein/lane) were separated by SDS-PAGE. Electrophoresis was conducted using a Xcell II mini cell with 4-12% NuPAGE Tris-acetate gradient SDS-polyacrylamide gels. Protein samples were transferred onto polyvinylidene difluoride membranes using an Xcell II Blot Module (Invitrogen). The membranes were blocked in Intercept® (TBS) blocking buffer (Licor Inc.) for 1 hour at room temperature, and probed with primary antibody in 0.1% Tween LiCor blocking buffer overnight at 4°C. The KLF14 antibody was a gift from Dr. Roger D. Cox (MRC Harwell Institute). It was generated at Eurogentec by immunizing rabbits against the C-DMIEYRGRRRTPRIDP-N peptide. Full length purified KLF14 purchased from Creative Biomart (Cat No. KLF14-481H) served as positive control. The β-actin antibody was from Cell Signaling Technology (Cat No. 3700s). The membranes were washed twice for 10 min each in TBS-0.01% Tween buffer solution and then probed with IR-Dye labeled secondary antibodies (926-32211 and 926-68072) in 0.1% Tween, 0.01% SDS LiCor blocking buffer for 1 hour at room temperature. Several washes in TBS-0.01% Tween buffer solution were repeated after labeling with secondary antibodies. The blots were scanned using the LiCor laser-based image detection method.

### Statistical analysis

Trial experiments were used to determine sample size with adequate statistical power. Measurement values that were beyond the boundary determined by the interquartile range were considered outliers and were excluded from statistical analyses. Since the scatter of the data follows normal distribution, all analyses were conducted with Student’s *t*-test with a two-tail distribution. Comparisons with *P* values <0.05 were considered significant. Effect size (β) was calculated by subtracting the mean difference between two groups, and dividing the result by the pooled standard deviation. β = (*M*_2_ - *M*_1_) ⁄ *SD*_pooled_, *SD*_pooled_ = √((*SD*_1_^2^ + *SD*_2_^2^) ⁄ 2); where M is the mean and SD is the standard deviation.

## RESULTS

### Adipocyte-specific deletion of KLF14 modifies adiposity, and insulin and lipid levels in female mice

Adipocyte-specific *Klf14* knockout (*Klf14*^fl/fl^*Adipoq*-Cre+ or KO) mice were born at expected Mendelian ratios and were indistinguishable from wildtype (*Klf14*^fl/fl^*Adipoq*-Cre- or WT) littermates at birth. At 8 weeks of age mice were placed on a 45% high fat (HF) diet (Research Diets D12079B). We measured body weights over a 12-week period (Figure 1A). Female *Klf14*^fl/fl^*Adipoq*-Cre+ mice showed similar body weight increases as WT littermates. However, male *Klf14*-deficient mice had lower body weight starting at 8 weeks on the HF diet (with 8% difference after 12 weeks on HF diet, *P*=0.018) (Figure 1B, left). In humans, lower KLF14 expression is associated with higher visceral fat mass (1). To determine whether loss of KLF14 in adipocytes affected adiposity on the HF diet, we measured body composition using EchoMRI (Figure 1A). We found increased fat mass relative to lean mass in female *Klf14*^fl/fl^*Adipoq*-Cre+ mice, while in male mice we observed the opposite (Figure 1B, middle and right). When determining individual organ weights normalized to tibia length, we found no differences between WT and *Klf14*-deficient mice for most organs, including brain, heart, kidney, muscle, spleen, and pancreas (Supplementary Figure 2). However, we observed that parametrial-periovarian fat depots in female *Klf14*-deficient mice were 70% larger (β_perigonadal_=1.105, *P*_perigonadal_=0.02), whereas subcutaneous and epididymal fat depots of male *Klf14*-deficient mice were 33% and 27% smaller compared to WT littermates (β_subq_=-1.073, *P*_subq_=0.01; β_epididymal_=-1.006, *P*_epididymal_=0.01) (Figure 1C). Notably, adipocyte *Klf14* deficiency resulted in decreased liver weight in males and a trend towards a decrease in females (β_male_=-0.626, *P*_male_=0.05; β_female_=-0.639, *P*_female_=0.18) (Figure 1C).

**Figure 1.**
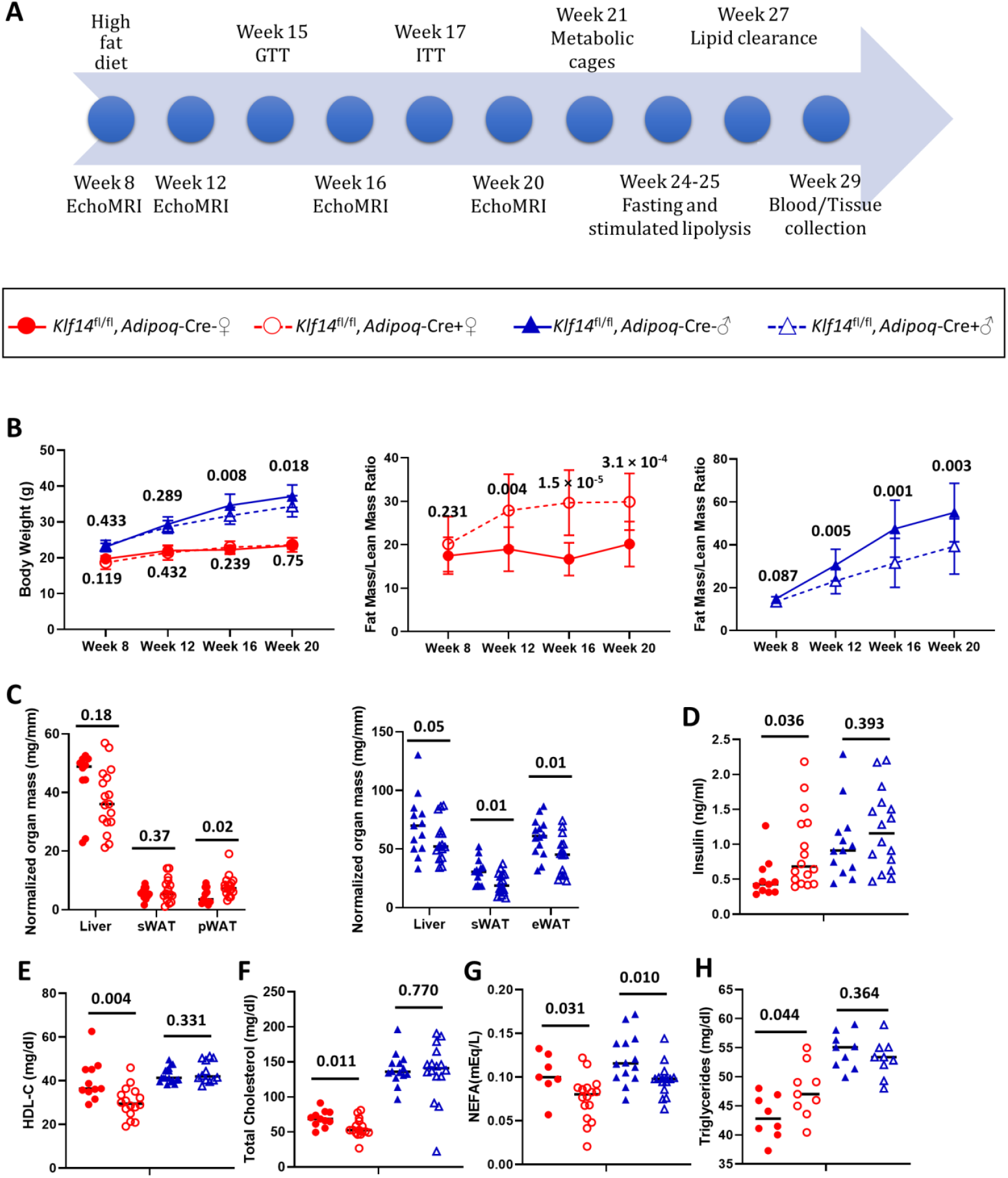
Adipocyte-specific deletion of KLF14 increases fat mass and affects circulating insulin and lipid levels in female mice. (**A)** Experimental design for the metabolic characterization of mice. (**B)** Body weight (left), fat mass/lean mass ratios in female (red) (middle, N_*Adipoq*-Cre-_ =11, N_*Adipoq*-Cre+_ =17) and male (blue) (right, N_*Adipoq*-Cre-_ =14, N_*Adipoq*-Cre+_ =14) mice at 8, 12, 16 and 20 weeks of age. (**C)** Normalized organ weights in females (left, N_*Adipoq*-Cre-_ =11, N_*Adipoq*-Cre+_ =17) and male mice (right, N_*Adipoq*-Cre-_ =14, N_*Adipoq*-Cre+_=14). sWAT denotes subcutaneous white adipose tissue; pWAT denotes parametrial-periovarian white adipose tissue; and eWAT denotes epididymal white adipose tissue. Plasma/serum levels of (**D)** insulin, (**E)** high-density lipoprotein C (HDL-C), **(F)** total cholesterol, **(G)** non-esterified free fatty acids (NEFA), and **(H)** triglycerides in female (N_*Adipoq*-Cre-_ =8-11, N_*Adipoq*-Cre+_ =9-17) and male (N_*Adipoq*-Cre-_ =9-14, N_*Adipoq*-Cre+_ =9-14) at 29 weeks of age after 6 hours of fasting. Data shown **(B)** are mean ∓ standard error of the mean and in **(C-H)** are mean and data for individual mice. *P*-values were calculated using two-tailed unpaired Student’s t -tests at each time point for (B) and all data in (C-H).

In humans, lower KLF14 expression in adipose tissue is associated with higher serum insulin and lower HDL levels with a stronger effect in females (1). Consistent with this, we observed 78% higher insulin and 25% lower HDL levels in *Klf14*-deficient female but not in male mice after a 6h fast (Figure 1D and E, β_female,insulin_=0.947, *P*_female,insulin_=0.036; β_female,HDL_=1.352, *P*_female,HDL_=0.004). Total cholesterol and non-esterified free fatty acids (NEFA) were lower by 19% and 26% while triglyceride (TG) levels were 10% higher in female mice with *Klf14* deficiency in adipocytes (Figure 1F, G and H, β_female,cholesterol_=1.176, *P*_female,cholesterol_=0.011; β_female,NEFA_=1.173, *P*_female,NEFA_=0.031, β_female,TG_=1.250, *P*_female,TG_=0.044). Mutant male mice also had lower NEFA levels (Figure 1G, β_male,NEFA_=1.022, *P*_male,NEFA_=0.01) but normal serum cholesterol and TG levels (Figure 1F and H).

### Adipocyte-specific deletion of KLF14 affects adipocyte size

The parametrial-periovarian fat depot in female mice and the epididymal fat depot in male mice are considered visceral fat. Larger visceral fat depots with bigger adipocytes are a significant risk factor for T2D, cardiovascular disease, and hypertension (24,25). Adipocyte size is increased in obese conditions (26,27). To test if adipocyte size was altered as a result of *Klf14* deficiency, we performed histological analysis using hematoxylin and eosin (H&E) staining of subcutaneous and perigonadal fat depots (Figure 2A). We observed that female mice with adipocyte deletion of *Klf14* had 25% smaller adipocytes in the subcutaneous depot (β_subq_=-0.422, *P*_subq_=4.02 × 10^−85^) and 70% larger adipocytes in the parametrial-periovarian depot relative to WT mice (β_periovarian_=0.706, *P*_periovarian_=2.25 × 10^−175^) (Figure 2B). In contrast, male mice with adipocyte deletion of *Klf14* had 6% larger adipocytes in the subcutaneous depot (β_subq_=0.09, *P*_subq_=1.66 × 10^−3^) and 14% smaller adipocytes in the epididymal depot relative to WT mice (β_epididymal_=-0.233, *P*_epidiymal_=2.66 × 10^−15^) (Figure 2B). These findings suggest that KLF14 regulates adipocyte size and adipose tissue mass in a depot specific and sex specific manner.

**Figure 2.**
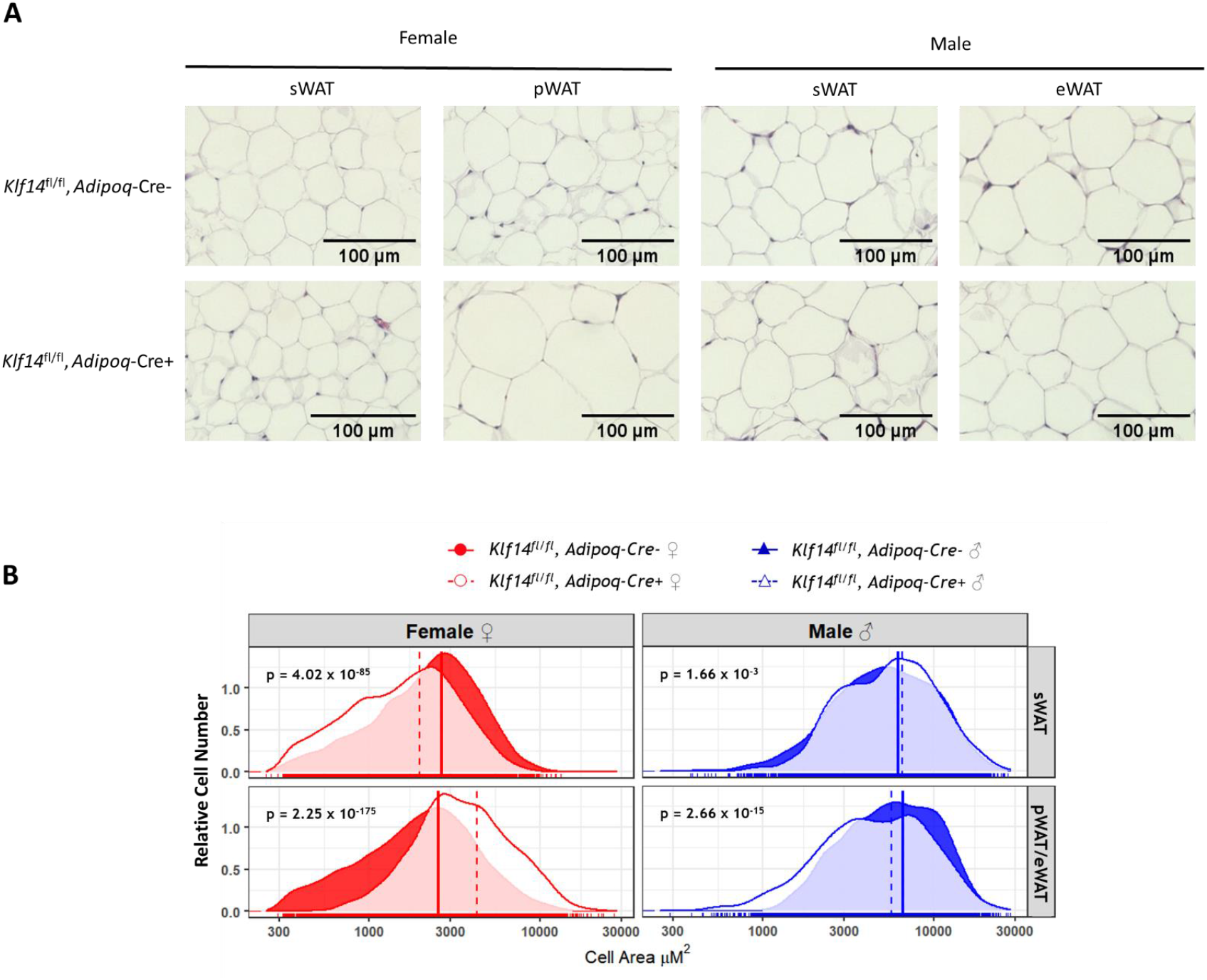
Adipocyte-specific deletion of KLF14 affects adipocyte size in female mice. (**A)** Representative subcutaneous (sWAT) and visceral (pWAT/eWAT) adipose tissue sections stained with Hematoxylin & Eosin. (**B)** Adipocyte size distribution in subcutaneous (sWAT) and visceral fat depots (pWAT/eWAT) {Citation}of female (red) and male (blue) mice. 5 µm-thick tissue sections from four mice for each genotype and sex were used for the analysis. The number of cells analyzed were as following: N_Female_*Adipoq*-Cre-_sWAT_ =3,988; N_Female_*Adipoq*-Cre+_sWAT_ =5,193; N_Female_*Adipoq*-Cre-_pWAT_ =4,132; N_Female_*Adipoq*-Cre+_pWAT_ =3,218; N_Male_*Adipoq*-Cre-_sWAT_ =2,699; N_Male_*Adipoq*-Cre+_sWAT_ =2,191; N_Male_*Adipoq*-Cre-_eWAT_ =2,196; N_Male_*Adipoq*-Cre+_eWAT_ =2,435 cells. *P*-values were calculated using two-tailed unpaired Student’s t -tests.

### Deletion of KLF14 in adipocytes causes increased insulin resistance in female mice

Adipose tissues regulate systemic glucose metabolism and insulin sensitivity (28,29). Genetic studies showed that T2D risk alleles associated with lower KLF14 expression in adipose tissue led to higher fasting insulin in humans indicating decreased insulin sensitivity (1,5). As described above, we also observed increased insulin levels in female *Klf14*-deficient mice (Figure 1D). To further explore whether the differences in adipose tissue mass as a result of *Klf14* deletion in adipocytes affected systemic glucose homeostasis and insulin sensitivity, we performed glucose and insulin tolerance tests with intraperitoneal injections of glucose or insulin after 7 and 9 weeks of HF diet, respectively. Mice with adipocyte-specific *Klf14* deletion had a similar response to an intraperitoneal glucose bolus as WT littermates (Figure 3A). However, in the insulin tolerance test female mice with adipocyte-specific *Klf14* deletion displayed increased glucose levels compared to WT littermates with 19% higher calculated area under the curve (AUC) (β_AUC_=1.233, *P*_AUC_=0.005) (Figure 3B) indicating increased insulin resistance. Insulin sensitivity for male mutant mice was similar to WT (β_AUC_=0.027, *P*_AUC_=0.943) (Figure 3B).

**Figure 3.**
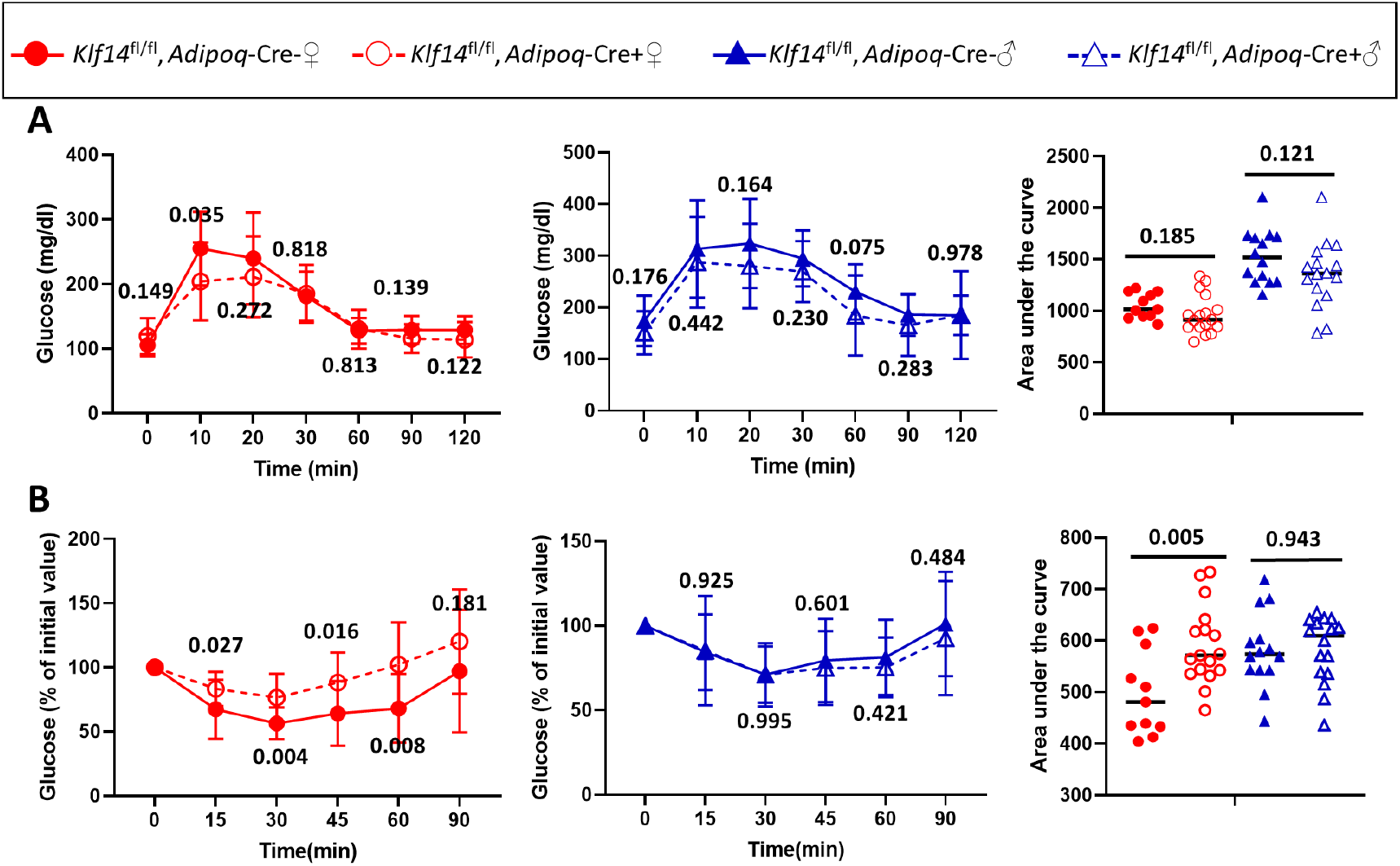
Adipocyte-specific deletion of KLF14 leads to increased insulin resistance in female mice. Blood glucose concentrations during intraperitoneal **(A)** glucose tolerance tests after 7 weeks of high fat diet and **(B)** insulin tolerance tests after 9 weeks of high fat diet in female (red; left) and male mice (blue; middle). Graphs for area under the curve (AUC) blood glucose concentrations are shown (right). N_Female_*Adipoq*-Cre-_ =11, N_Female_*Adipoq*-Cre+_ =17, N_Male_*Adipoq*-Cre-_ =14, N_Male_*Adipoq*-Cre+_ =14. Data are mean ∓ standard error of mean. *P*-values were calculated using two-tailed unpaired Student’s t -tests at each time point.

### Deletion of KLF14 in adipocytes modifies energy metabolism in female mice

Increased fat mass in female mice with *Kl14* deficiency suggested a defect in energy balance. To investigate the role of KLF14 in energy homeostasis, we performed indirect calorimetry studies using metabolic cages. Food consumption was comparable between *Klf14*-deficient and WT mice (Supplementary Figure 4), and mice of both genotypes maintained body weights within 5-10% of their initial weight during the three-day period in the metabolic cages. Despite these similarities, female *Klf14*^fl/fl^*Adipoq*-Cre+ mice showed a 38% decrease in locomotor activity (β_Locomotor activity_=2.212, *P*_Locomotor activity_=5.2 × 10^−4^) compared to WT mice during the dark cycle (Figure 4A). Oxygen consumption (VO_2_) was lower in both the light and dark cycles by 26% and 19%, respectively (β_O2, dark_=2.320, *P*_O2, dark_=0.003; β_O2, light_=1.569, *P*_O2, light_=0.0095) while CO_2_ release was 14% lower in the light cycle and trended towards a decrease in the dark cycle (β_CO2, dark_=0.900, *P*_CO2, dark_=0.093; β_CO2, light_=1.262, *P*_CO2, light_=0.026) (Figures 4B and C). The calculated respiratory exchange ratio 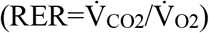, an indicator for the metabolic energy source (carbohydrate or fat), was higher (close to 1) for female *Klf14*^fl/fl^*Adipoq*-Cre+ mice than for WT littermates (β_RER,dark_=4.258, *P*_RER,dark_=1.0 × 10^−6^) in the dark cycle (Figure 4D), suggesting predominant use of carbohydrates as energy source, despite being fed a HF diet. The indirect calorimetry measurements allowed us to estimate consumed substrates using gas exchange ratios as a surrogate for substrate utilization (16). Female *Klf14*^fl/fl^*Adipoq*-Cre+ mice had 82% higher carbohydrate utilization (β_carbohydate,dark_=2.015, *P*_carbohydrate,dark_=0.01) and 79% decreased fat utilization (β_fat,dark_=-5.400, *P*_fat,dark_=1.1 × 10^−6^) in the dark cycle compared to WT mice. We did not observe differences for any of the measured parameters for male mice in dark or light cycles (Figure 4).

**Figure 4:**
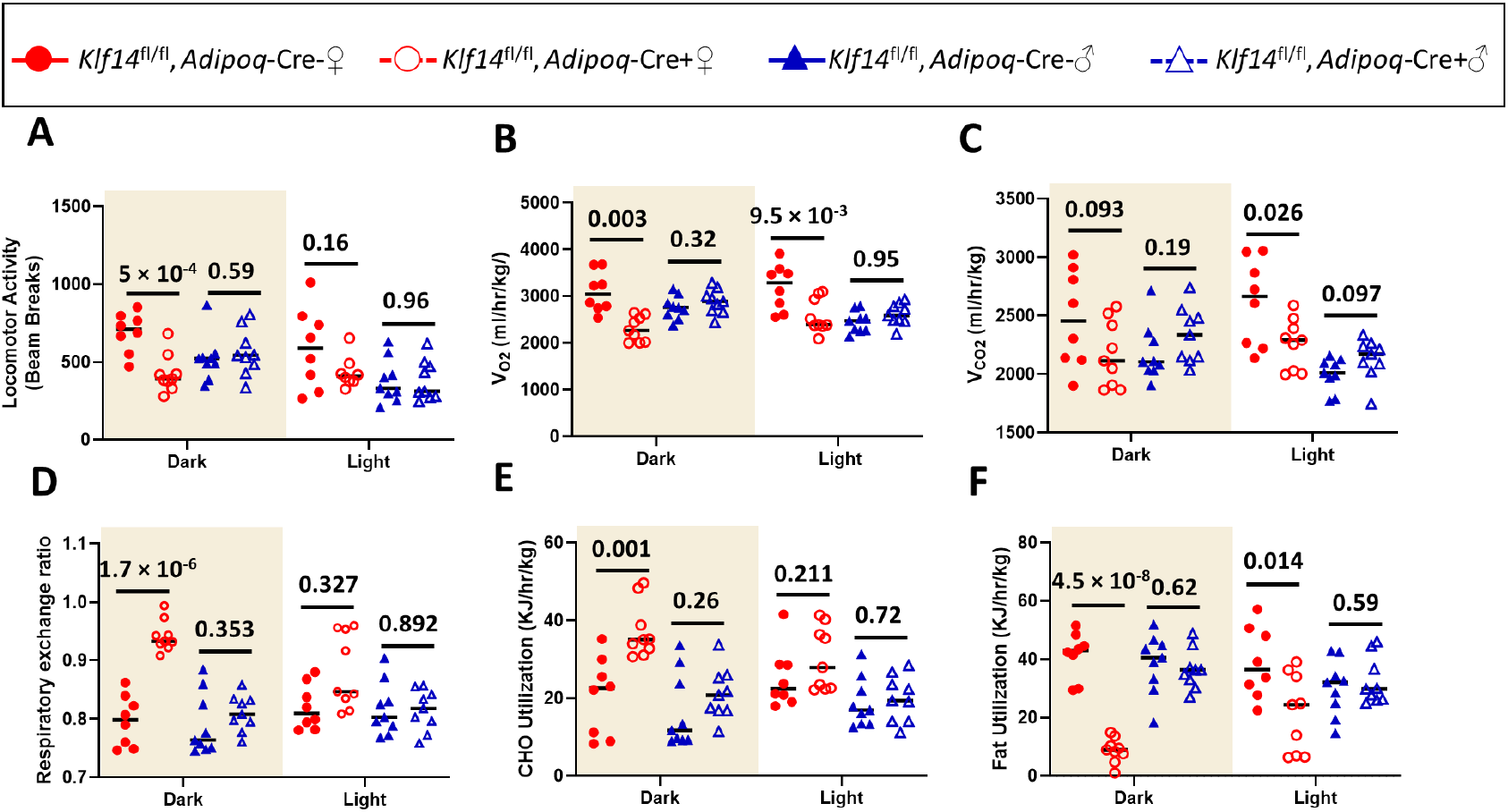
Adipocyte-specific deletion of KLF14 modifies energy metabolism in female mice. Indirect calorimetry studies measuring **(A)** total activity, **(B)** oxygen consumption (VO2), **(C)** carbon dioxide production (VCO_2_), **(D)** respiratory exchange ratio, **(E)** carbohydrate (CHO) utilization, and **(F)** fat utilization in female (red) and male (blue) mice during dark (shaded area) and light cycles. N_Female_*Adipoq*-Cre-_ =8, N_Female_*Adipoq*-Cre+_ =9, N_Male_*Adipoq*-Cre-_ =9, N_Male_*Adipoq*-Cre+_ =9. Mean and individual mouse data are shown. *P*-values were calculated using two-tailed unpaired Student’s t -tests.

### Deletion of KLF14 in adipocytes causes defects in lipid metabolism in female mice

We reasoned that defects in lipid breakdown (lipolysis) or lipid uptake and synthesis (lipogenesis) in adipocytes could explain the shift in energy source for female mice. Compatible with these possibilities, circulating NEFA and triglyceride levels were decreased and increased under fasting conditions, respectively, in female mutant mice (Figures 1G and H). To follow up on these changes, we first measured lipolytic products, non-esterified fatty acids (NEFA) and glycerol, after 16 hours of fasting. NEFA levels increased as a result of fasting, as expected (Figure 5A). However, female *Klf14*^fl/fl^*Adipoq*-Cre+ mice had 17% and 38% lower NEFA levels compared to WT littermates in fed and fasted states (β_fed_=-1.416, *P*_fed_=0.013; β_fasted_=-3.231, *P*_fasted_=1 × 10^−4^) (Figure 5A) with NEFA levels in *Klf14*-deficient female mice increasing to a lesser extent compared to WT littermates in response to fasting (β=-1.292, *P*=0.021) (Figure 5B). Glycerol in female mice was only decreased in the fed state (β_fed_=2.157, *P*_fed_=7 × 10^−4^) (Figure 5C and D). No differences in NEFA and glycerol were observed in male mice except that NEFA under fed conditions were slightly increased (p=0.036) (Figure 5).

**Figure 5:**
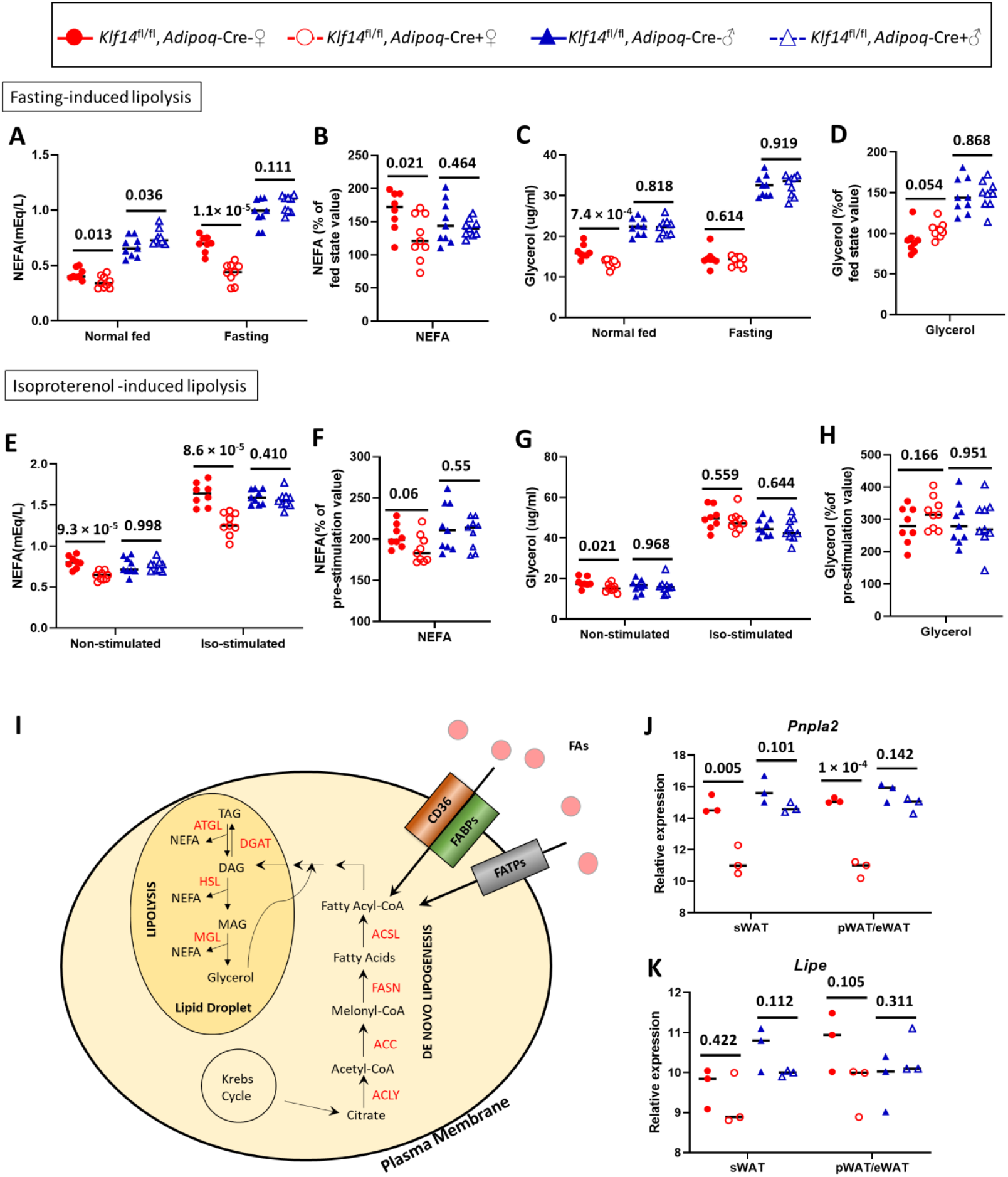
Adipocyte-specific deletion of KLF14 causes defects in lipolysis in female mice. Mice after a 16-week high-fat diet were fasted for 16 hours. **(A and B)** Fasting non-esterified free fatty acids (NEFA) and **(C and D)** glycerol were measured and compared to that of mice before fasting. Mice after a 17-week high fat diet were treated with 10 mg/kg isoproterenol. Plasma **(E and F)** NEFA and **(G and H)** glycerol was measured and compared to that of mice before stimulation. N_Female_*Adipoq*-Cre-_ =8, N_Female_*Adipoq*-Cre+_ =9, N_Male_*Adipoq*-Cre-_ =9, N_Male_*Adipoq*-Cre+_ =9. Changes in plasma levels of NEFA, and glycerol were calculated relative to baseline values (fed-state for B and D or before stimulation for F and H). Mean and individual mouse data are shown. *P*-values were calculated using two-tailed unpaired Student’s t-tests. Data from female mice are shown in red and male mice are shown in blue. **(I)** Schematic representation of genes playing a role in triglyceride breakdown (lipolysis) and formation (lipogenesis), and fatty acid uptake. Lipolysis is catalyzed by three lipases: adipose triglyceride lipase (ATGL), hormone-sensitive lipase (HSL), and monoacylglycerol lipase (MGL) whose actions result in the release of NEFA and glycerol. Triglyceride synthesis through de novo lipogenesis uses citrate to make acetyl-CoA by ATP-citrate lyase (ACL), acetyl-CoA is converted to malonyl-CoA by acetyl-CoA carboxylase (ACC), and fatty acid synthesis is achieved by conversion of malonyl-CoA into fatty acids by fatty acid synthase (FASN). Fatty acids are then made into fatty acyl-CoAs by acyl-CoA synthetase (ACSL) and converted to diacylglycerols (DAG) through several enzymatic reactions. Diacylglycerol acyltransferase (DGAT) then catalyzes the conversion of DAG into TAG (triacylglycerol or triglyceride). Cellular uptake of fatty acids (FAs) is facilitated by CD36, FABPs and FATPs. mRNA expression of **(J)** *Pnpla2* and **(K)** *Lipe* in mature adipocytes isolated from subcutaneous (sWAT) and parametrial-periovarian/epididymal WAT (pWAT/eWAT). N =3 mice per genotype and sex. Relative gene expression, normalized to GAPDH levels, was calculated using the 2^-ΔΔCT^ method. *P*-values were calculated using two-tailed unpaired Student’s t -tests.

Adipocyte lipolysis is regulated by hormonal and neuronal stimulation (30) with beta-adrenergic receptor signaling potently increasing lipolysis (31). To further corroborate that female adipocyte *Klf14*-deficient mice indeed had a defect in lipolysis, we thus quantified serum levels of NEFA and glycerol in response to isoproterenol that stimulates beta-1 and beta-2 adrenergic receptors (32).Female *Klf14*^fl/fl^*Adipoq*-Cre+ mice had 21% and 22% lower NEFA levels compared to WT littermates under fed condition in non-stimulated and stimulated states, respectively (β_non-stimulation_=-2.692, *P*_non-stimulation_= 9 × 10^−5^; β_stimulated_=-2.699, *P*_stimulated_ = 8 × 10^−5^) (Figure 5E) with NEFA levels in *Klf14*-deficient female mice increasing to a lesser extent than in WT littermates in response to stimulation (β=-1.022, *P*=0.06) (Figure 5F). No differences were observed in male mice (Figure 5E and F). Similar to fasting experiments, we observed a difference in serum glycerol level in female mice before stimulation (β_female_=-0.307, *P*_female_ = 0.021) but not after (β_female_=-0.301, *P*_female_ = 0.559) (Figure 5G). Changes in serum glycerol levels upon stimulation did not differ between genotypes in both female or male mice (β_female_=0.815, *P*_female_ = 0.166; β_male_=-0.031, *P*_male_ = 0.951) (Figure 5H). The difference in NEFA and glycerol responses to induced lipolysis is most likely due to different metabolic fates of NEFA and glycerol after release into circulation.

TG hydrolysis to NEFA and glycerol is achieved in a three-step enzymatic pathway (Figure 5I). We measured the expression of adipocyte triglyceride lipase (ATGL), the first enzyme in the TG breakdown pathway and found that mRNA levels of *Pnpla2* in adipocytes isolated from subcutaneous and parametrial-periovarian/epididymal fat were decreased by 24% and 29%, respectively, in female mice as a result of *Klf14* deficiency (β_sWAT_=-9.320, *P*_sWAT_=5 × 10^−3^; β_pWAT_=-13.139, *P*_pWAT_=1 × 10^−4^) (Figure 5J), but were normal in male mice. We also measured hormone sensitive lipase (HSL), the second enzyme in the TG breakdown pathway, but did not observe differences in *Lipe* mRNA expression in adipocytes of male or female mice (Figure 5K), which may be due to the fact that HSL activity is predominantly regulated by phosphorylation (33). Lower *Pnpla2* expression is consistent with above-described decreased lipolysis and increased fat mass and adipocyte size of visceral adipocytes in female mutant mice. Visceral adipocytes are normally more lipolytically active when compared to subcutaneous adipocytes (34).

Our metabolic cage data indicated that female *Klf14*^fl/fl^*Adipoq*-Cre+ mice were not able to efficiently use dietary fatty acids leading us to speculate that there was also a defect in lipid uptake. Therefore, we challenged mice with a bolus of olive oil by oral gavage and measured TG and NEFA levels to quantify lipid clearance from the bloodstream. We observed that blood TG and NEFA clearance were both decreased in female mutant mice compared to WT mice (β_4hr-TG_=4.610, *P*_4hr-TG_=7.2 × 10^−9^; β_4hr-NEFA_=3.528, *P*_4hr-NEFA_=8 × 10^−6^) (Figure 6A and B), consistent with decreased use of lipids as an energy source in female *Klf14*-deficient mice. No differences were observed in male mice (Figure 6C and D). These results are consistent with our metabolic cage data that female mice use less fat as their energy source.

**Figure 6:**
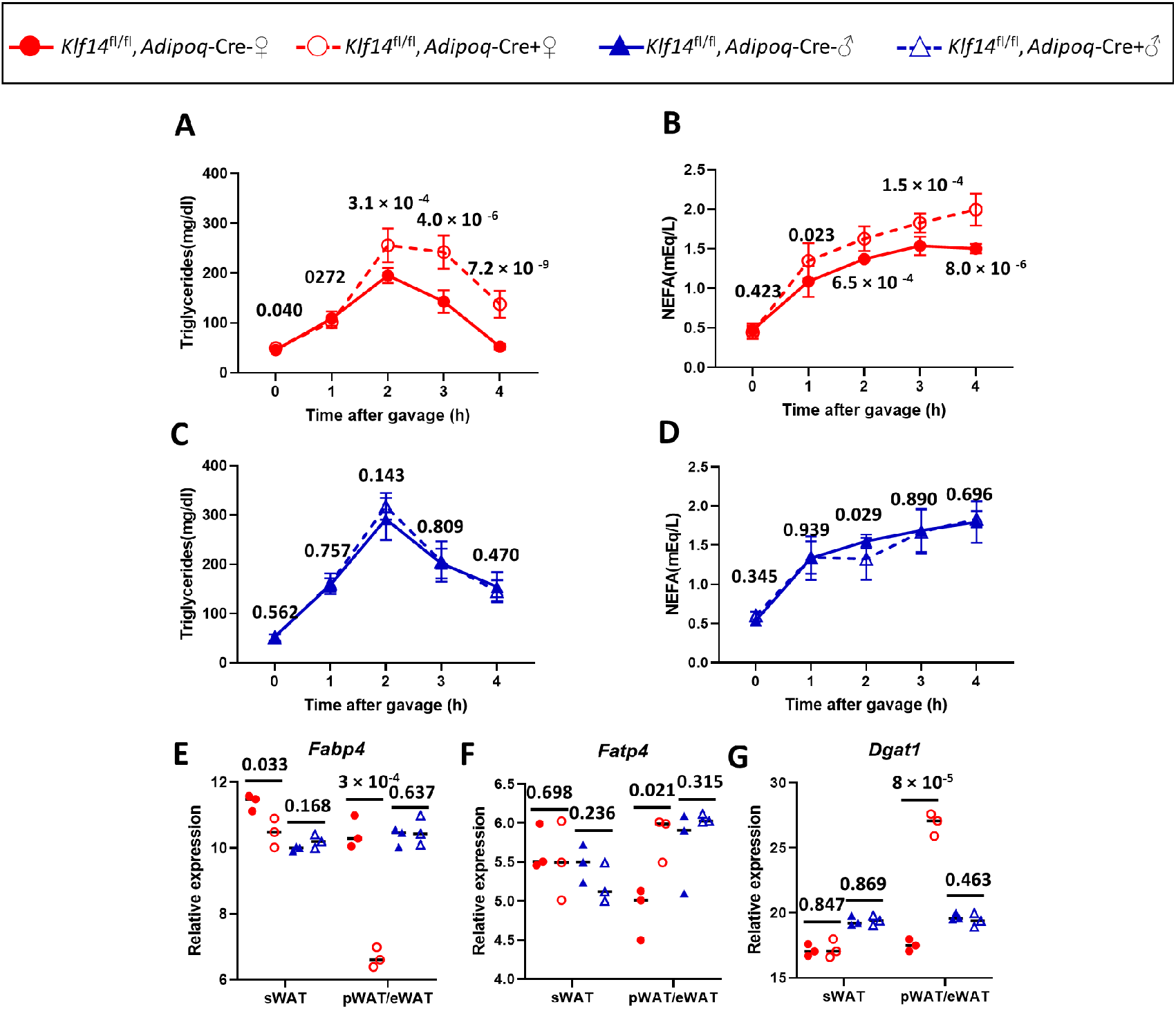
Adipocyte-specific deletion of KLF14 decreases lipid clearance in female mice. Mice on a 19-week high-fat diet were gavaged with a bolus of olive oil. Plasma triglycerides in **(A)** female and **(C)** male mice and non-esterified free fatty acids (NEFA) in **(B)** female and **(D)** male mice were measured over four hours. N_Female_*Adipoq*-Cre-_ =8, N_Female_*Adipoq*-Cre+_ =9, N_Male_*Adipoq*-Cre-_ =9, N_Male_*Adipoq*-Cre+_ =9. mRNA expression of fatty acid uptake gene **(E)** *Fabp4* **(F)** *Fatp4* and de novo lipogenesis gene **(G)** *Dgat1* were measured in mature adipocytes isolated from subcutaneous (sWAT) and parametrial-periovarian/epididymal WAT (pWAT/eWAT). N =3 mice per genotype and sex. Relative gene expression, normalized to GAPDH levels, was calculated using the 2^-ΔΔCT^ method. Mean and standard error of the mean are shown in A, B, C, D; mean and individual mouse data are shown in E, F and G. *P*-values were calculated using two-tailed unpaired Student’s t -tests. Data from female mice are shown in red and data for male mice are shown in blue.

Uptake of fatty acids into cells is achieved predominantly via a saturable protein-facilitated process (35) (Figure 5I). Several proteins facilitate the uptake of fatty acids, among which the best characterized are CD36, FABPs and FATPs (36–38). Fatty acids are also synthesized in cells through *de novo* lipogenesis that is catalyzed by ATP-citrate lyase (encoded by *Acly*) (39), acetyl-CoA carboxylase (encoded by *Acaca* and *Acacb*) (40), and fatty acid synthase (encoded by *Fasn*) (41). Fatty acids are converted into fatty acyl-CoAs by acyl-CoA synthetase (encoded by *Acsl*) and then esterified with glycerol to form TGs, with the last step catalyzed by diglyceride acyltransferase (encoded by *Dgat*) (42). We assayed several key enzymes in adipocyte fatty acid uptake and lipogenesis and observed sex- and depot-specific differences in *Fabp4, Fatp4* and *Dgat1* mRNA expression in isolated mature adipocytes. *Fabp4* was 36% lower in the parametrial-periovarian depot of female *Klf14*-deficient mice (β_Female pWAT_=-10.750, *P*_Female pWAT_=3 × 10^−4^) (Figure 6E) while *Fatp4* (Figure 6F) and *Dgat1* (Figure 6G) was 20% and 54% higher in the parametrial-periovarian depot of the same mice compared to WT controls (β_Female pWAT_=14.336, *P*_Female pWAT_=8 × 10^−5^). We also observed differences in other pathway genes that were either sex- or depot-specific. mRNA levels for *Fatp1* were decreased in both female subcutaneous WAT and perigonadal/visceral WAT, *Dgat2* was decreased in male subcutaneous WAT and epididymal/visceral WAT. We also observed that Fabp5 was decreased in both female and male subcutaneous WAT and visceral WAT (Supplementary Figure S4). *Acaca, Acacb* and *Fasn* showed no differences. (Supplementary Figure S4).

### Adipocyte KLF14 overexpression reduces body fat in female mice

Our results showed that KLF14 deficiency resulted in an adverse metabolic phenotype in female mice. We therefore hypothesized that overexpression of *Klf14* may be metabolically beneficial. To address this, we generated a transgenic mouse with *Klf14* overexpression under the regulation of the *Adipoq* promoter (*Adipoq-Klf14-*OE) (see Material and Methods section for a detailed description). This resulted in 1.7-fold induction of KLF14 in adipocytes (Supplementary Figure 1C). We fed these mice a high fat diet for 18 weeks and performed body composition analysis using EchoMRI. Male and female mice from both genotypes gained similar amounts of body weight, and no difference in body weight between the *Klf*14-overexpressing mice (*Adipoq-Klf14-*OE) and WT littermates were observed at the end of the 18-week diet challenge (Figure 7A). However, starting at 12 weeks of HFD, female *Adipoq-Klf14-*OE mice showed a trend towards decreased fat mass to lean mass ratios compared to WT littermates (Figure 7B, *P*=0.074). At the end of the study, after 18 weeks of high fat diet, female mice with *Klf14* overexpression in adipocytes had 38% lower body fat compared to WT littermates (Figure 7B, β_Female_=-1.446, *P*_Female_= 0.002). No differences were observed in male mice with *Klf14* overexpression (Figure 7C, P=0.99).

**Figure 7:**
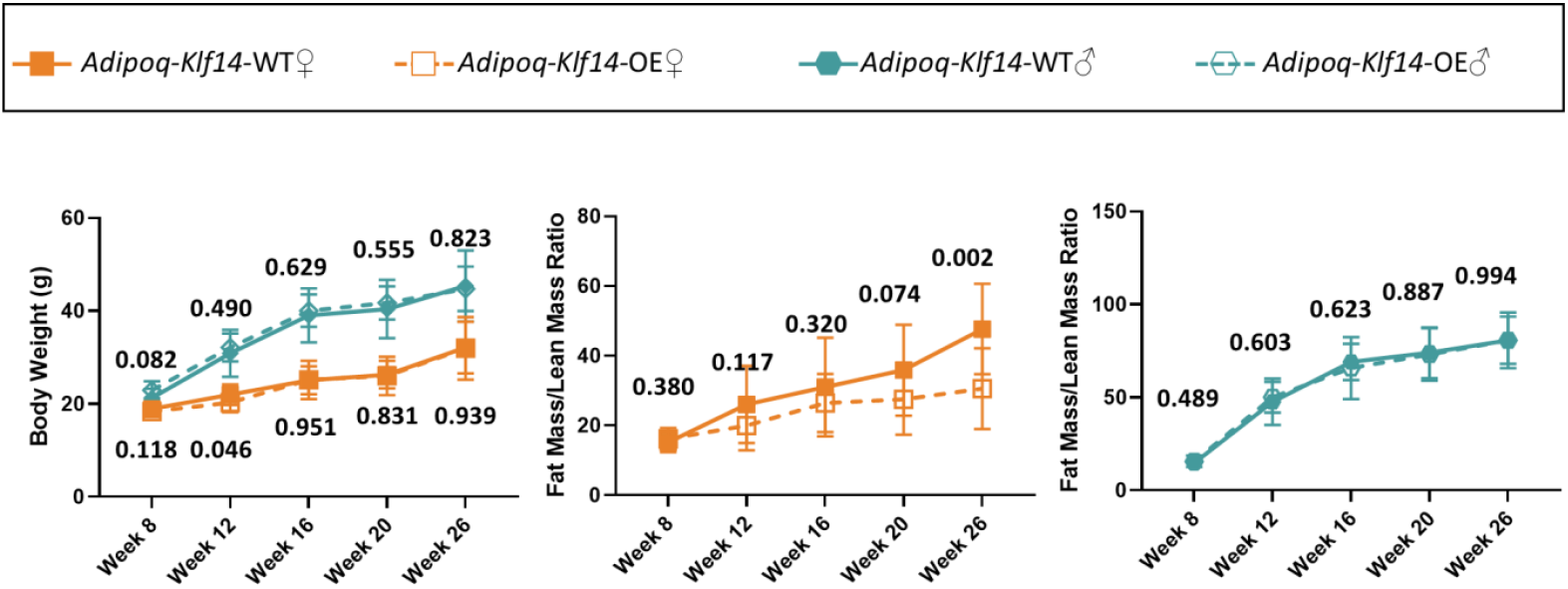
Adipocyte-specific overexpression of KLF14 decreases fat mass in female mice. Body weight (left), fat mass/lean mass ratio in female (orange) (middle, N_WT_ =14, N_*Adipoq*-OE_ =14) and male (blue) (right, N_WT_ =8, N_*Adipoq*-OE_ =12) mice at 8, 12, 16, 20, and 26 weeks of age. Data shown are mean ∓ standard error of the mean. *P*-values were calculated using two-tailed unpaired Student’s t -test at each time point.

In summary (Figure 8), *Klf14* deficiency in adipocytes caused major abnormalities in high fat diet-fed female but not male mice. Female adipocyte *Klf14*-deficient mice had increased total body fat with larger visceral fat mass and bigger visceral adipocytes, but they had smaller subcutaneous adipocytes. Concomitant with the changes in body fat distribution and adipocyte size, the mice demonstrate metabolic abnormalities including insulin resistance and altered serum lipid levels. Despite high fat diet feeding, *Kfl14*-deficient female mice used carbohydrates as the predominant source of energy and simultaneously had lower energy expenditure and activity. Further evaluation of lipid metabolism uncovered defects in fasting- and beta-adrenergic induced lipolysis, and impaired triglyceride and NEFA clearance after lipid gavage. These defects were coupled with decreased mRNA expression of *Pnpla2* (a key lipolytic enzyme) in both perigonadal/visceral and subcutaneous adipocytes. Furthermore, mRNA levels for proteins involved in cellular fatty acid uptake including Fabp4, Fabp5, and Fatp1 were decreased in both subcutaneous WAT and perigonadal/visceral WAT, but mRNA levels of the fatty acid transport protein Fatp4 and Dgat1, an enzyme that catalyzes the conversion of DAG into triglycerides, were increased only in perigonadal/visceral adipocytes. In contrast, adipocyte-*Klf14*-deficient male mice on a HFD gained less body weight, had decreased total body fat with smaller epididymal/visceral and subcutaneous fat depots and adipocytes, but had, relative to wild type, normal metabolic parameters and insulin sensitivity. The male also did not show defects in stimulated lipolysis and lipid clearance, and mRNA levels of enzymes involved in cellular lipid metabolism were not different from wild type littermates, except that Fatp1 was increased in sWAT, and Fabp5 and Dgat2 decreased in subcutaneous WAT and epididymal/visceral WAT. In conclusion, our findings suggest that KLF14 regulates adipose tissue mass and adipocyte size in a depot specific and sex specific manner most likely by differentially modifying adipocyte lipid metabolism.

**Figure 8:**
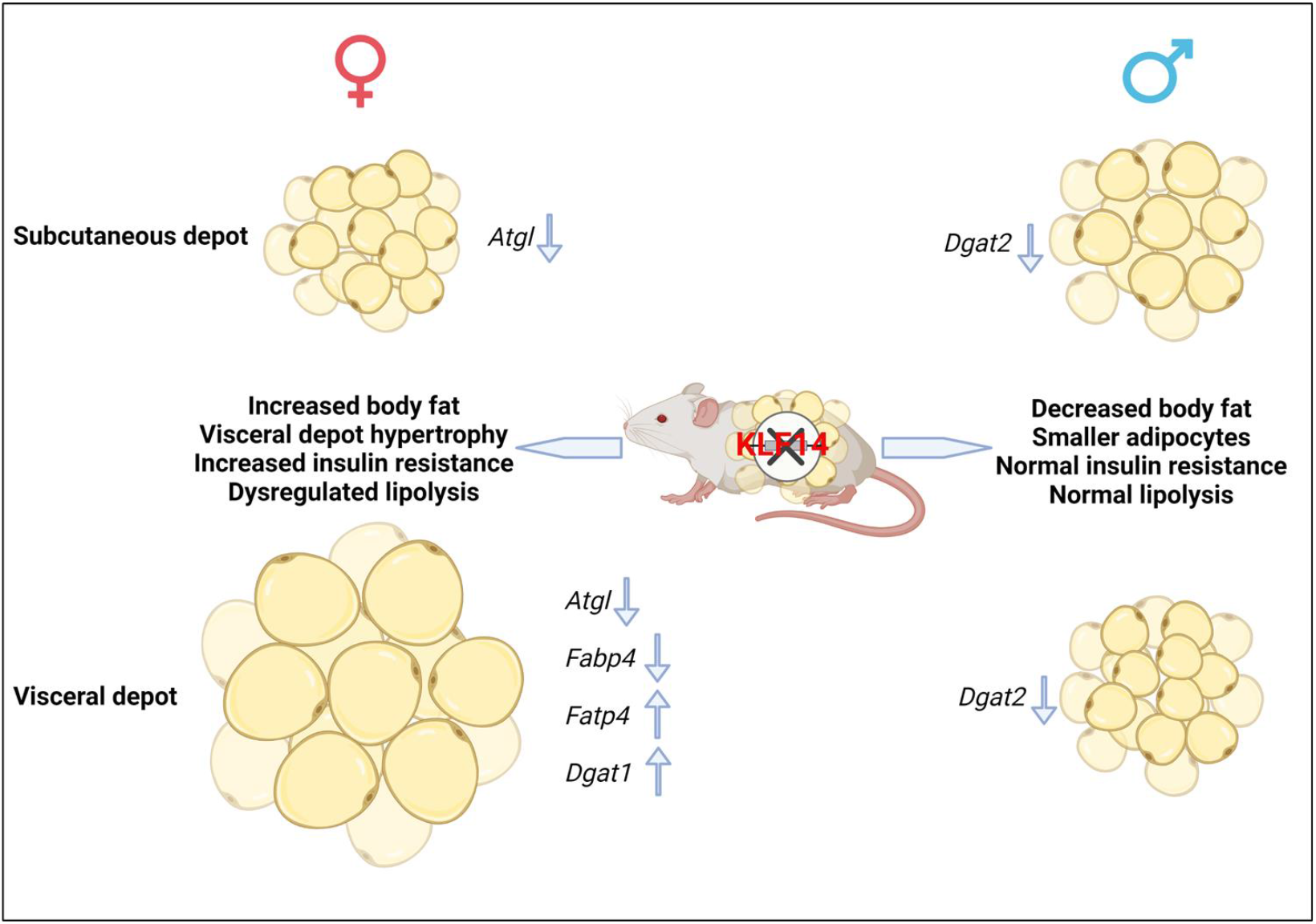
Summary of main findings. Deletion of KLF14 in mouse adipocytes resulted in sex-dimorphic and depot-specific differences. Female mice with adipocyte *Klf14*-deficiency had higher total body fat, primarily due to increased visceral depot fat mass. Female mutant mice also displayed a shift in body fat storage from subcutaneous depot to visceral depot as evidenced by visceral adipocyte hypertrophy and decreased adipocyte size in subcutaneous depots. Meanwhile, male mice had lower body fat mass and smaller cell size in the visceral depots. Female *Klf1*4 knockout mice were insulin resistant while insulin sensitivity in male mice with adipocyte *Klf14*-deficiency was not different from wildtype littermates. Changes in adipose tissue mass and adipocyte size can, at least in part, be explained by dysregulation of lipid metabolism, including lipolysis, fatty acid uptake and lipogenesis. Several genes involved in these pathways are dysregulated in a sex- and depot-specific manner as indicated.

## DISCUSSION

Visceral adipose tissue expansion independent of overall adiposity is associated with an increased risk for developing T2D and cardiovascular disease (43). How deposition of excess calories into the different adipose depots is regulated is currently unknown. However, sex-dependent differences in adipose tissue distribution are well-documented with higher adiposity in females than males (44). Males usually display “android” or “apple shape” distribution with more visceral fat in the abdominal region, while females have “gynoid” or “pear shape” distribution with more subcutaneous fat and less visceral fat in the lower body (45). A shift in fat deposition from subcutaneous depots around the hip to visceral depots in the waist region in males confers increased risk for metabolic disorders (46). However, females can harbor increased upper-body fat without enhanced metabolic disease risk as excess fat is mainly stored in subcutaneous depots. Our results suggest that the transcription factor KLF14 plays a role in modifying adipocyte function and thereby adipose tissue distribution in a sex- and depot-specific manner. Deletion of the *Klf14* gene in mature adipocytes resulted in an increase of visceral fat in female mice but a decrease in male mice. The shift in lipid storage from subcutaneous to parametrial-periovarian (visceral) depots in female mice with adipocyte *Klf14* deletion was characterized by changes in adipocyte size, smaller adipocytes in subcutaneous and larger adipocytes in perigonadal/visceral adipose tissues and major abnormalities in lipid metabolism.

Sex hormones play a significant role in sex-specific body fat distribution, as evidenced by the fact that android and gynoid fat distribution patterns in males and females appear as early as puberty (47). However, circulating levels of testosterone, progesterone and estradiol at 29 weeks of HFD were comparable between female and male adipocyte-*Klf14*-deficient and wild type mice (Supplementary Figure 5), suggesting that the sex differences we observed in mice are likely hormone independent. It is however possible that differential expression of sex hormone receptors or sex-specific interactions of KLF14 with sex hormone receptors could play a role in transcriptional target expression; other KLF family transcription factors cobind DNA with the estrogen receptor. We do not know how KLF14 affects body fat distribution and impacts adipocyte function distinctly in different sexes. KLF14 has been shown to be a master regulator of adipose tissue gene expression (1). We speculate that differential expression of sex hormone receptors or sex-specific interactions of KLF14 with sex hormone receptors could play a role in transcriptional target expression, especially since, other KLF family transcription factors have been shown to cobind DNA with the estrogen receptor (48). Primary targets in male and female adipocytes need to be established.

The absence of *Klf14* in adipocytes causes significant abnormalities in lipid metabolism, both lipolysis and fatty acid uptake in female mice. Whether these are primary consequences of KLF14 deficiency or secondary metabolic abnormalities due to changes in adipose tissue mass and distribution, and adipocyte size is not entirely clear. Future identification of the primary consequences of KLF14 deficiency on metabolism and primary Klf14 targets will require the careful analysis of metabolic phenotypes at various time points before and after the transition to HF diet. Abnormalities in circulating lipids in female mice with *Klf14*-deficiency in adipocytes can partly be explained by the increased insulin resistance in adipocyte *Klf14*-deficient female mice. This includes the lower circulating HDL-C and increased plasma triglyceride levels under fasting condition, and impaired triglyceride and NEFA clearance after lipid gavage. Triglyceride clearance requires the action of lipoprotein lipase in adipose tissues and insulin-stimulated lipoprotein lipase activity is impaired in insulin resistance. Insulin-stimulated NEFA uptake under fed conditions is also defective in insulin resistance. Furthermore, expression of fatty acid transporters is decreased in the presence of impaired insulin sensitivity (49). However, the lower-than-normal NEFA levels in female mice under fed and fasting conditions and in response to beta-adrenergic stimulation cannot be attributed to impaired insulin sensitivity. On the contrary, NEFA levels should be increased due to defective suppression of lipolysis by insulin. Currently, we do not know the mechanisms by which *Klf14*-deficiency in adipocytes impairs lipolysis. But impaired lipolysis is accompanied by a decrease in *Pnpla2* mRNA in adipocytes of both sWAT and pWAT in female mice. Since KLF14’s DNA binding motif is not well-established, future studies will establish if ATGL is a primary target of KLF14.

Hormone sensitive lipase (HSL) was once thought to be the key enzyme that regulates cellular TG breakdown. Its activity is modulated by phosphorylation (33) However, *Hsl* deficient mice had a normal rate of FFA production, indicating that other enzymes are playing a key role in TG breakdown (50). Haemmerle *et al*. identified ATGL as the rate limiting enzyme in the mobilization of cellular TG (51). Notably, they reported that mice with inactivated ATGL displayed deleterious metabolic phenotypes similar to those of our female adipocyte *Klf14-*deficient mice. These include increased adipose tissue mass, decreased insulin sensitivity, and decreased availability of FFAs (51). The latter fostered the use of carbohydrates as the primary fuel despite the presence of increased amounts of fat in the adipose tissue. Reduced TG hydrolysis then led to reduced energy expenditure and total activity. This study supports the notion that defective lipolysis in adipocyte *Klf14*-deficient mice leads to the switch to carbohydrates as the predominant source of energy, the decrease in energy expenditure and activity, and increased fat mass gain. The fact that glucose tolerance is not impaired in adipocyte *Klf14*-deficient mice despite impaired insulin sensitivity may be explained by the energy substrate switch to glucose and decreased cellular NEFA uptake facilitating instead glucose uptake into tissues.

The increased size of parametrial/visceral adipocytes in females with adipocyte *Klf14*-deficiency can be explained by impaired lipolysis as visceral adipocytes are normally more lipolytic than subcutaneous adipocytes (34). But increased expression of *Dgat1* in parametrial/visceral adipocytes may further contribute to increased adipocyte size. Mice overexpressing Dgat1 in adipocytes have increased deposition of TG in WAT (52). Although fatty acid uptake is impaired in adipocytes, as decreased expression of key fatty acid binding/transporter proteins and impaired NEFA clearance suggest, fatty acids taken up could be efficiently esterified to TG due to the increase in *Dgat1*. Increased *Dgat1* expression cannot be explained by increased insulin resistance. In contrast, Dgat1 expression positively correlates with insulin sensitivity in human subjects (53). Specific transcription factors regulating Dgat1 expression have not been identified, but PPARγ is a candidate (54). Whether KLF14 directly or indirectly controls Dgat1 expression will need to be investigated.

Interestingly, male mice despite decreased fat mass and smaller adipocytes did not show improved metabolic parameters and greater insulin sensitivity when compared to wild type littermates. It is possible that simultaneous changes in fatty acid transporter/binding proteins in the smaller adipocytes are leading to unfavorable changes that are, however, offset by improved insulin sensitivity.

Previous mouse studies showed conflicting results for KLF14 on metabolic phenotypes and atherosclerosis. We recently reviewed these studies (5), several of which only used male mice to study the impact of KLF14 deletion or ectopic overexpression, a consequential omission given that in humans and in our study, KLF14 plays sexual-dimorphic roles. Some of the studies also used different diet compositions which may be the source of the conflicting results.

Finally, we showed a metabolically favorable consequence of increased expression of adipocyte KLF14 since female overexpression mice accumulated less body fat in response to a HF diet challenge. Previous studies showed that perhexiline, an approved therapeutic small molecule presently in clinical use to treat angina and heart failure, induced KLF14 expression and reduced atherosclerosis (55). Our results, together with the results from this previous pharmacological study, suggest that therapeutic targeting of KLF14, specifically in adipose tissue, may lead to an improved metabolic phenotype.

## Acknowledgements

We thank the members of the Civelek laboratory for their feedback and discussion. We thank Dr. Stephen B. Abbott for the assistance in indirect calorimetry analysis. We thank Dr. Michael M. Scott for the assistance in EchoMRI.

## Author Contributions

Q. Yang and M. Civelek conceived the study; J. Hinkle, J.N. Reed, R. Aherrahrou, Z. Xu participated in data collection and analysis. K. Musunuru generated *Klf14*^fl/fl^*Adipoq*-Cre+ mice. T. E. Harris, E. J. Stephenson, and S.R. Keller contributed to the experimental design and data interpretation. Q. Yang, S.R. Keller and M. Civelek drafted the article. M. Civelek directed the study. All authors edited the final article. M. Civelek is the guarantor of this work and, as such, had full access to all the data in the study and takes responsibility for the integrity of the data and the accuracy of the data analysis.

## SUPPLEMENTAL FIGURE LEGENDS

**Supplemental Figure 1:**
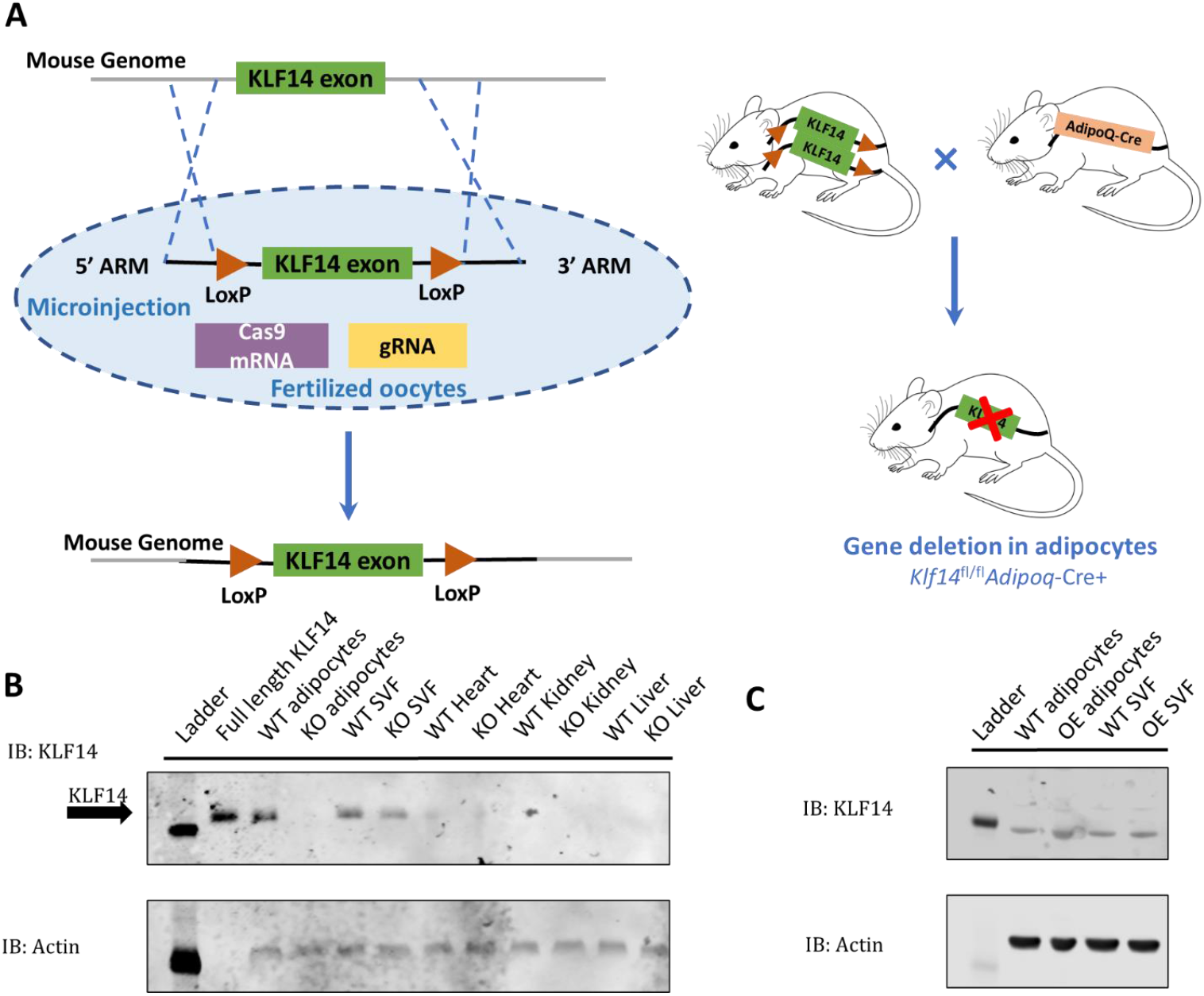
Characterization of transgenic mice.**(A)** Targeting strategy for deletion of Klf14 in mouse adipocytes. **(B)** KLF14 protein level in adipocytes, stromal vascular fraction (SVF), heart, kidney and liver of wild-type (WT) and knockout (KO) mice. **(C)** KLF14 protein level in adipocytes, stromal vascular fraction (SVF) of wild-type (WT) and overexpression (OE) mice.

**Supplemental Figure 2:**
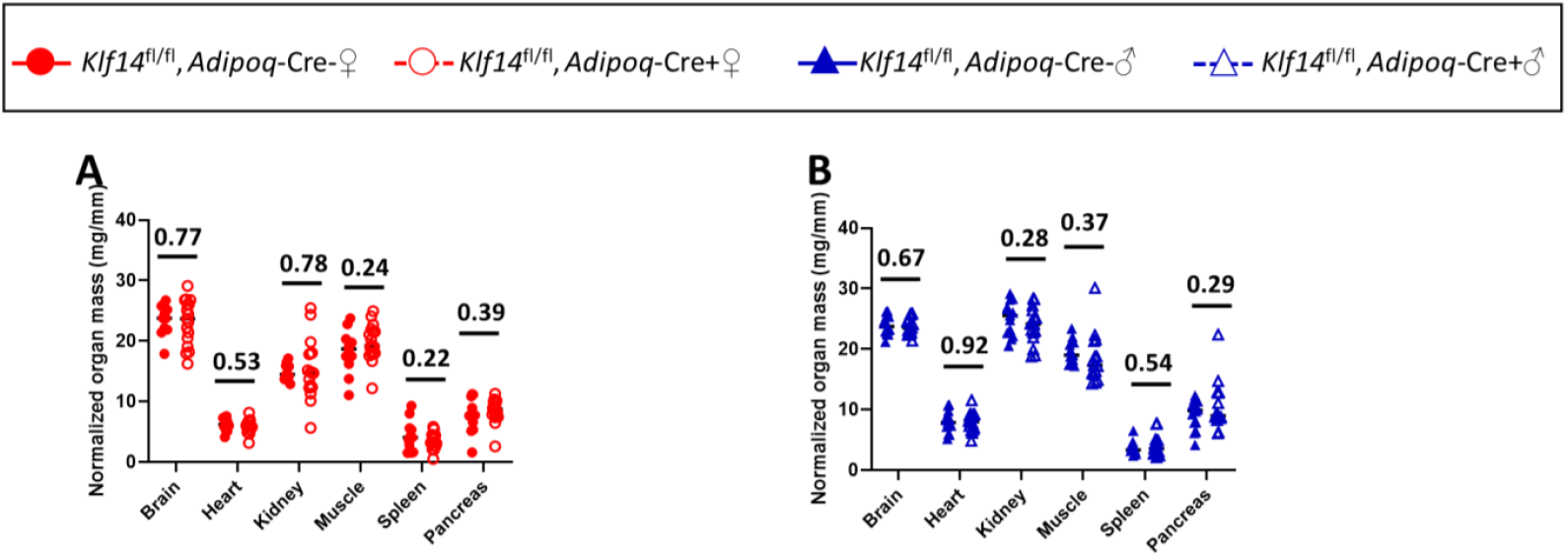
Tissue weights normalized to tibia length in adipocyte-specific *Klf14* knockout and wild type **(A)** female and **(B)** male mice. N_Female, *Adipoq*-Cre-_ =11, N_Female,A*dipoq*-Cre+_ =17, N_Male,*Adipoq*-Cre-_ =14, N_Male,*Adipoq*-Cre+_=14.

**Supplemental Figure 3:**
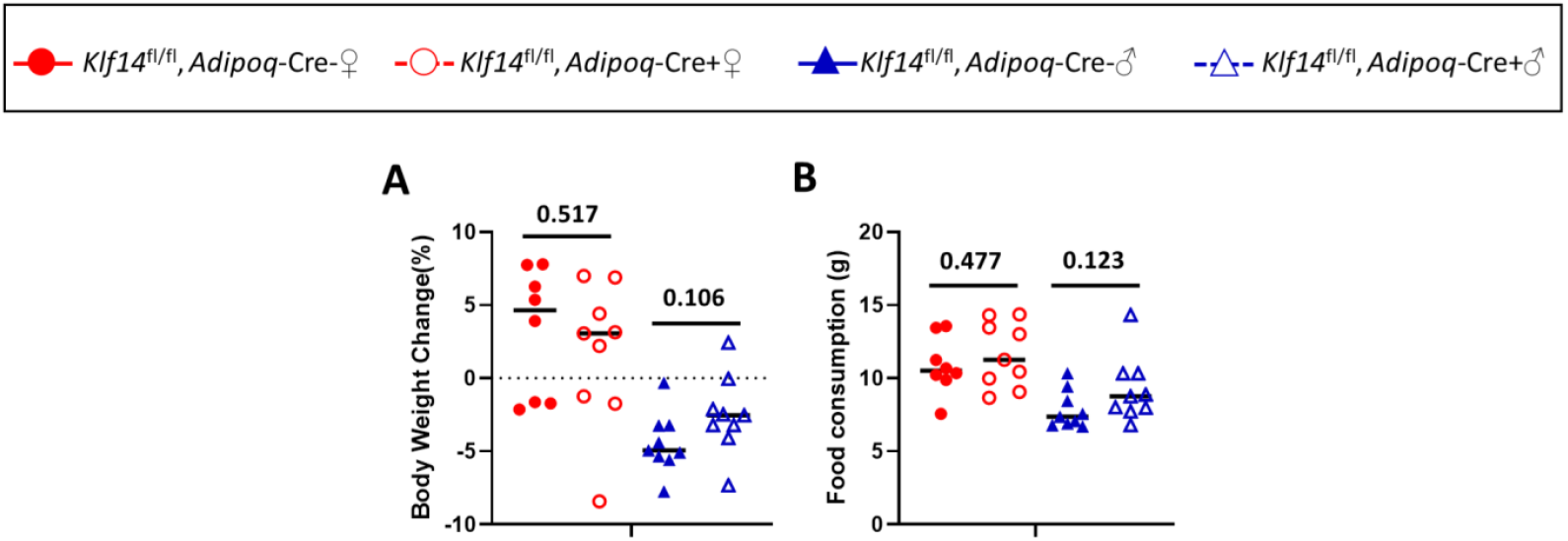
**(A)** Body weight change and **(B)** food consumption during the three-day period in metabolic cages. N_Female_*Adipoq*-Cre-_ =8, N_Female_*Adipoq*-Cre+_ =9, N_Male_*Adipoq*-Cre-_ =9, N_Male_*Adipoq*-Cre+_ =9.

**Supplemental Figure 4:**
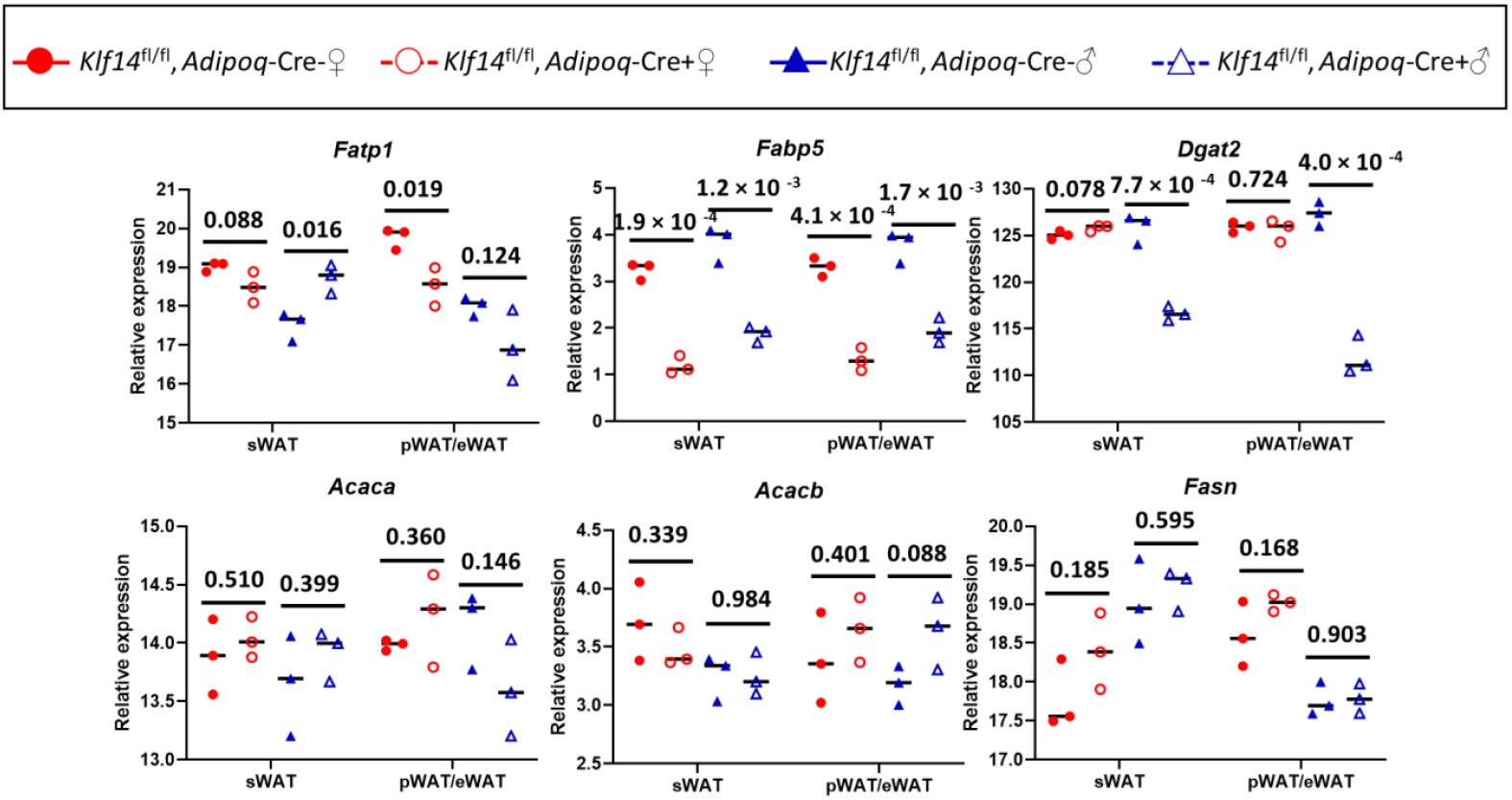
mRNA expression of fatty acid uptake and metabolism genes *Fatp1, Fabp5, Dgat2, Acaca, Acacb, Fasn* and in isolated mature adipocytes. N =3 mice per genotype and sex. Relative gene expression, normalized to GAPDH levels, was calculated using the 2^-ΔΔCT^ method. *P*-values were calculated using two-tailed unpaired Student’s t -test.

**Supplemental Figure 5:**
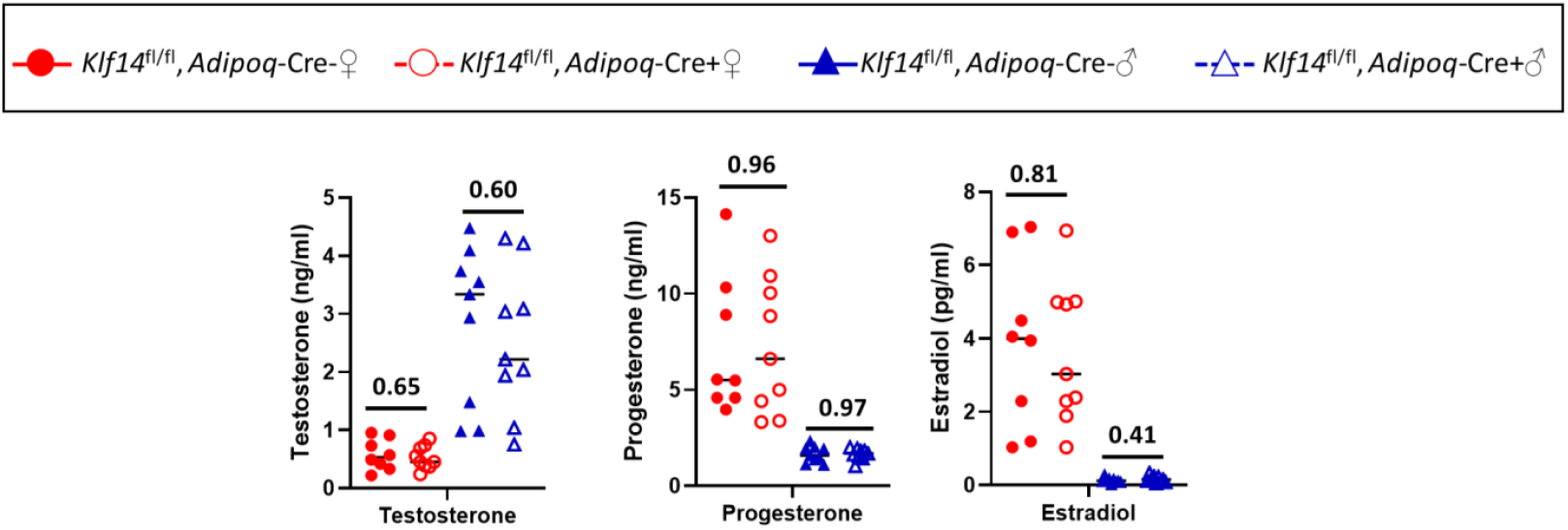
Sex hormone levels in serum of adipocyte *Klf14*-deficient female and male mice and control littermates at euthanasia after 21 weeks of HFD. (A) Testosterone, (B) Progesterone, (C) Estradiol. N_Female_*Adipoq*-Cre-_ =8, N_Female_*Adipoq*-Cre+_ =9, N_Male_*Adipoq*-Cre-_ =9, N_Male_*Adipoq*-Cre+_ =9.

**Supplemental table 1:**
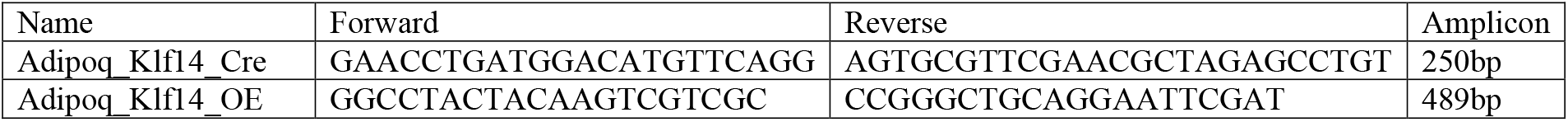
Mouse genotyping primer list.

**Supplemental table 2:**
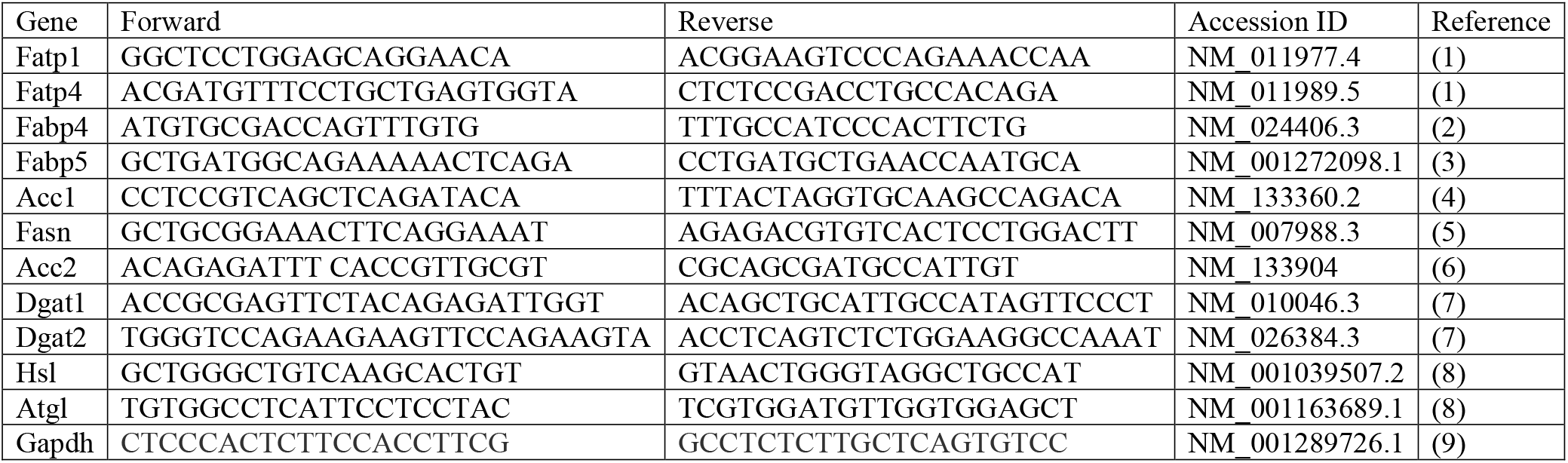
qPCR primer list.

